# SMIM36, a novel and conserved microprotein, is involved in retinal lamination in zebrafish

**DOI:** 10.1101/2023.01.14.524032

**Authors:** Surbhi Sharma, Soundhar Ramasamy, Yasmeen Khan, Dheeraj Chandra Joshi, Beena Pillai

## Abstract

Microproteins are small proteins comprising 2 to 200 amino acids, arising from small Open Reading Frames (smORFs). They are found in different parts of the cell and regulate basic molecular processes like DNA replication, repair, transcription and recombination. SMIM or SMall Integral Membrane proteins are novel, largely uncharacterized, members to the class of microproteins defined by the presence of a transmembrane domain. The retinal transcriptome of zebrafish, reported previously by our group, revealed several novel mRNA transcripts that show oscillating expression in a diurnal manner. Here, we show that one of these transcripts encodes the zebrafish homolog of the human SMIM36 protein, which has not been functionally characterised. This highly conserved microprotein is expressed in the human and zebrafish retina, and efficiently translated in cell lines. Using single-cell RNA-seq datasets, we found that it is expressed in the bipolar cells, rods and Muller glia in the human retina. The knockdown of SMIM36 using splice-block morpholino caused microphthalmia and defects in the retinal layers in zebrafish. Therefore, we show the role of a microprotein in the neural retina thus paving the way for future studies on the role of SMIM proteins in retinal disorders.

## Introduction

The completion of the human genome project has led to the identification of a large number of genes of unknown function, a significant fraction of which are thought to be non-coding. In parallel, attempts to characterise the translatome comprehensively have revealed the presence of a substantial number of short ORFs that could potentially give rise to proteins. Assuming that shorter ORFs will not form stable protein products, initially, ORF sequences of <100 amino acids were not considered biologically relevant. With recent advancements in ribosome profiling methods, mass spectrometry and algorithms to analyse the resulting data, a large number of peptides arising from non-canonical ORFs are being detected. These ORFs and peptides have been referred to variously, based on their size (microproteins), origin (smORFs, altORFs), structure (cyclotides) and often inhibitory role (miPs, ID proteins)(Jiang, Lou, and Hou 2021).

Microproteins are defined as small proteins comprising 2 to 200 amino acids, arising from small Open Reading Frames (smORFs). They are widely found and frequently overlooked but can be involved in a variety of biological phenomena like development, regeneration and transcriptional regulation. At the molecular level, some microproteins called miPs act by interfering in multi-protein complexes and dimers. However, the large number of uncharacterized microproteins suggest that new molecular mechanisms may be revealed as more of these enigmatic proteins are studied. The functional relevance of a number of small proteins like DWORF (Q. Zhang et al. 2017; Makarewich et al. 2018), Mitoregulin (Stein et al. 2018) and Minion (Q. Zhang et al. 2017) in muscle performance, PIGBOS (Q. Zhang et al. 2017) and BRAWNIN (S. Zhang et al. 2020) in ER and mitochondria function and NoBody (D’Lima et al. 2017) in mRNA post-transcriptional regulation, has been revealed recently. Although in vivo studies are rare, certain microproteins have been implicated in disease related phenotypes. For instance, depletion of PINT87aa, a candidate tumor suppressor, leads to rapid proliferation of cancer cells and over-expression showed a reduction in the tumorigenic potential in vivo(M. Zhang et al. 2018).

SMIMs or SMall Integral Membrane proteins are a recent addition to the growing list of microproteins. These are small proteins containing approximately 100 amino acids conserved across vertebrates. They comprise a family of unrelated proteins with expression in diverse cellular and subcellular compartments, similarly named owing to the presence of a single-pass transmembrane domain. The human protein atlas provides information on sub-cellular address of various SMIM proteins like cytoplasm (SMIM20, 22, 30), mitochondria (SMIM4, 8, 12) and nuclear speckles (SMIM19) (Thul et al. 2017). Most of them remain uncharacterized with the exception of SMIM1 (D’Lima et al. 2017; Arnaud et al. 2015) in blood cells and SMIM30 (Pang et al. 2020) in hepatocellular carcinoma.

In a screen for transcripts that show circadian rhythmicity, we identified a transcript called pc10 (Ramasamy et al. 2019). This transcript contains a small coding region which could potentially encode a microprotein of 102 aa length. On the basis of conservation, the microprotein shows homology to the uncharacterized SMIM36 microprotein in humans. In this study, we demonstrate the biological relevance of SMIM36 in the neural retina. The retina, a highly networked neural tissue present at the back of the eye, is vital for visual function. SMIM36 expression was originally detected in the zebrafish retina, as part of a previous study by our group, focusing on genes that are expressed diurnally in the retina (Ramasamy et al. 2019). Single-cell RNA-seq (scRNA-seq) analysis of publicly available datasets revealed SMIM36 expression in the bipolar cells of both zebrafish and human retina, with expression in Muller glia and rods in the human retina. The gene co-expression analysis of the scRNA-seq dataset of larval zebrafish revealed the co-expression with bipolar cell-specific transcription factors further strengthening the cell-type specific expression. Using morpholino mediated knockdown in zebrafish embryos we show that loss of SMIM36 leads to microphthalmia and retinal lamination defects. Thereby, our work provides the first evidence of occurrence and function of a conserved SMIM protein in the neural retina, the loss of which leads to retinal defects, paving the way for future work on deciphering the role of these proteins in retinal anomalies of humans.

## Results

### SMIM36, a novel and conserved microprotein localized to cytoplasm is expressed in teleost and human retina

In zebrafish, the gene ENSDARG00000105527.2 is associated with a 884nt RNA transcript which contains a 309nt long coding region that can be translated into a 102 amino acid long protein, which is 57% conserved to the SMIM36 protein predicted from the human gene (ENSG00000261873). Both proteins contain a highly conserved, central transmembrane domain flanked by relatively less conserved N and C-terminal domains (Figure1A). In spite of 90% of the mRNA transcript comprising non-coding regions, the ability of the transcript to code for a protein is supported by various lines of evidence. Firstly, it was detected in the mass spectrometry profile of the human retina (Kim et al. 2014), proving it to be a bonafide protein-coding gene in the retina. The evidence of its translation is also corroborated by ribosome profiling in developing zebrafish embryos (Bazzini et al. 2014); (Michel et al. 2014). We cloned the SMIM36 coding region as an in-frame covalent fusion with eGFP and confirmed that it is translated and expressed stably in mammalian cell lines (Figure 1B) where it showed a cytoplasmic localization.

**Figure 1.**
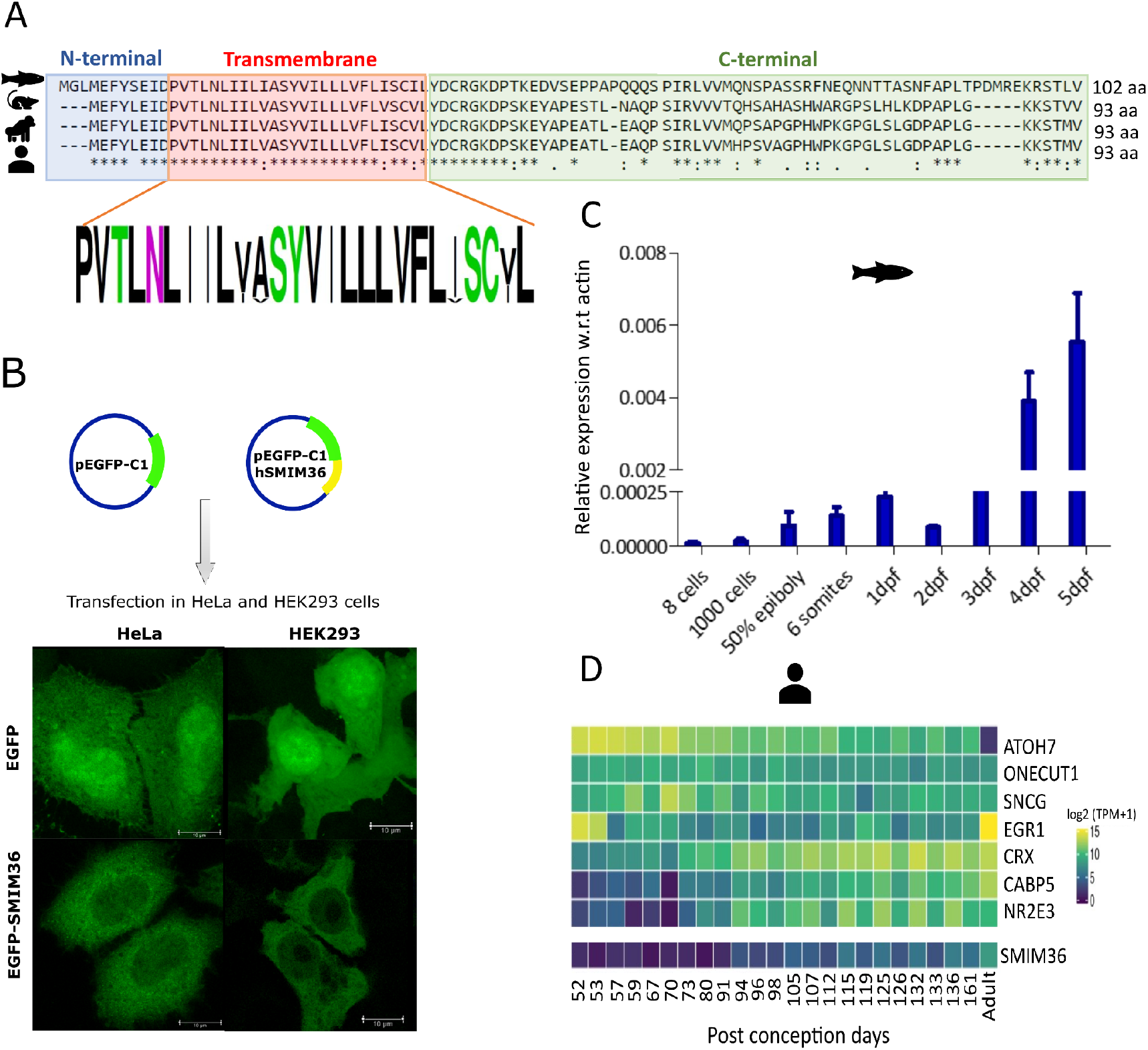
SMIM36, a novel and conserved microprotein is expressed in teleost and human retina. A) Multiple sequence alignment of SMIM36 protein sequence across vertebrate species with logo plot highlighting the conserved transmembrane domain. B) Schematic showing plasmid constructs used for expressing EGFP and EGFP-SMIM36 in HeLa and HEK293 cell lines. Fluorescent images showing the localization of EGFP-SMIM36 protein tagged with EGFP to cytoplasm. scale bar- 10μm. C) Real- time quantification of SMIM36 RNA during developmental stages of zebrafish. Bars represent mean±s.e.m. across five biological replicates, dpf-days post fertilization. D) Expression profile of SMIM36 and retinal cell type marker genes across developing human retina (Heatmap generated by Eye in a Disk online database).

Next we studied the expression profile of SMIM36 mRNA in zebrafish development. During zebrafish development, its expression is detected first at 50% epiboly stage (Figure 1C), a stage when neural development commences (Woo and Fraser 1995). In publicly available RNA-seq datasets of the human retina, we observed its expression at 94th day post-conception (Figure 1D) (Swamy and McGaughey 2019).

### scRNA-seq reveals SMIM36 expression in bipolar cells, rods and Muller glia in the human retina

The cell-type specific expression of a novel gene is an important clue to its probable function, especially in the retina since it contains a highly heterogeneous mix of cells, intricately arranged in their respective layers. Broadly, the outer nuclear layer contains photoreceptors, followed by the inner nuclear layer containing interneurons like bipolar cells and the projection layer containing retinal ganglion cells. To assign the cell-type identity to SMIM36, we utilised publicly available scRNA-seq datasets of the retina. In zebrafish (Farnsworth, Saunders, and Miller 2020) larvae, it showed expression in the bipolar cells of the inner nuclear layer of the retina in agreement with our own observations from in situ hybridization (Ramasamy et al. 2019). For the human retina, we re-analyzed the scRNA-seq dataset of Lukowski et al. (Lukowski et al. 2019) using the Seurat package (Satija et al. 2015) (Figure 2A). The human retinal cells formed 15 distinct cell clusters (Figure 2B), in which SMIM36 expression was distributed in various cell clusters (Figure 2C). To dissect the identity of SMIM36 expressing cells, we subsetted the count data of SMIM36 positive cells and performed Seurat clustering (Figure 2D). Further cluster identification using known retinal markers revealed SMIM36 expression in bipolar cells, rods and Muller glia of the human retina (Figure 2E). Overall, our analysis revealed that in addition to sequence conservation, the cell-type specific expression of SMIM36 is also conserved across vertebrates.

**Figure 2.**
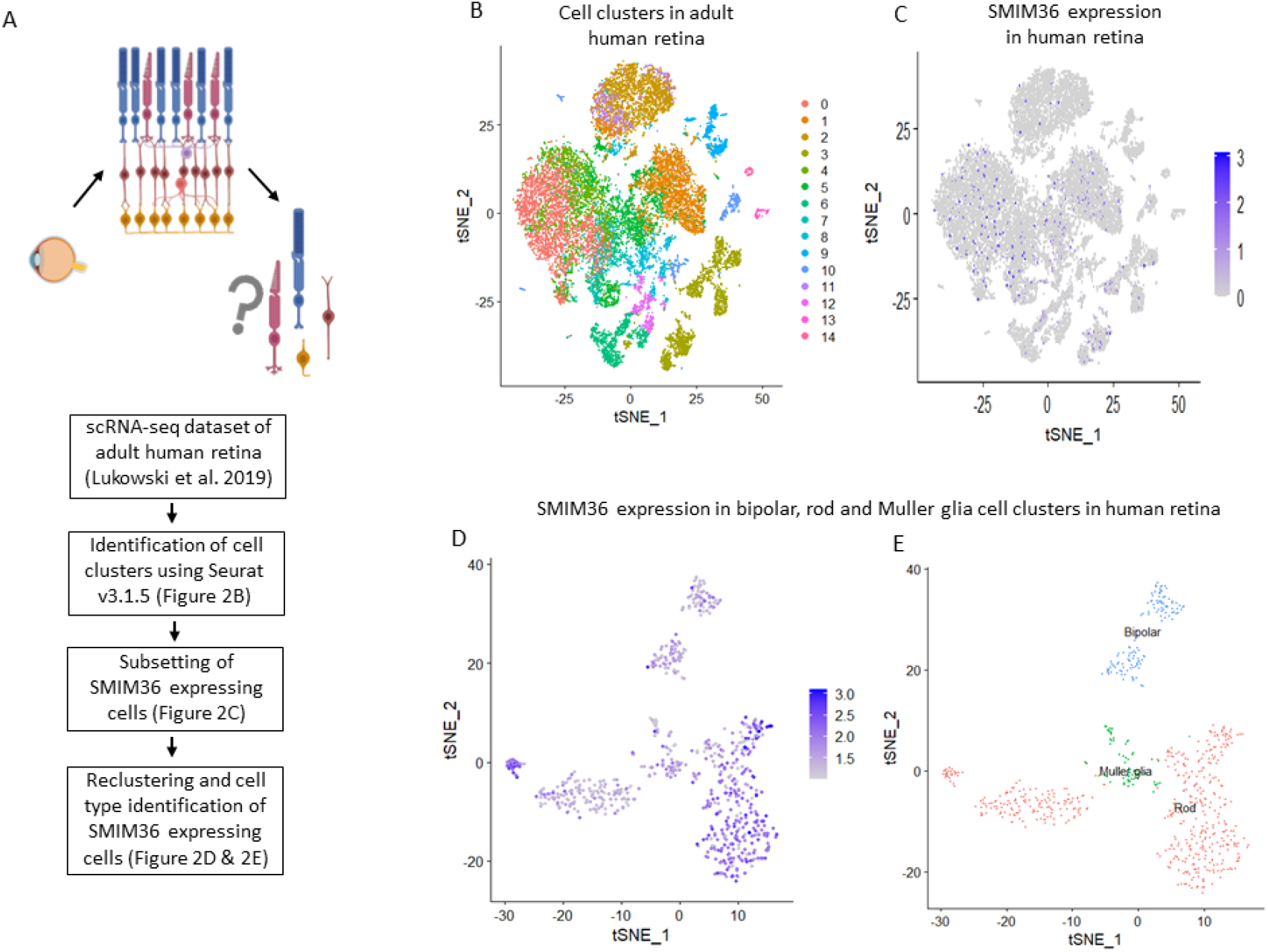
SMIM36 is expressed in bipolar cells, rods and Muller glia in the human retina. A) Schematic of the workflow used for the analysis of scRNA-seq dataset of adult human retina (dataset source-Lukowski et al. 2019). B) t-SNE plot showing cell clusters in the scRNA-seq dataset of adult human retina (n=3). C) Feature plot showing SMIM36 expression in various clusters derived from human retina. D) Feature plot obtained after subsetting and reclustering of SMIM36 expressing cells. E) Identification of cell clusters (Figure 3D) using known markers of retinal cell types.

### Co-expression analysis reveals potential role for SMIM36 in eye development

Often genes involved in a particular biological process show correlated expression patterns, a feature of the activity of transcription factors. The bipolar cell-specific expression of SMIM36 suggested its regulation by a cell-type specific transcription factor. To investigate this, we used GRN-Boost in the SCENIC (Single-Cell rEgulatory Network Inference and Clustering) analysis workflow (Aibar et al. 2017), which identifies co-expressing gene pairs or modules.

SCENIC inputs a list of transcription factors (TFs) and a single-cell gene expression matrix to identify co-expressing TFs and their target genes (Figure 3A). For this study, we subsetted the single-cell gene expression matrix of SMIM36 expressing cells derived from scRNA-seq dataset of 5dpf zebrafish larvae (Spanjaard et al. 2018) (FigureS2A&B). GRN-Boost analysis on the above dataset revealed strong co-expression of SMIM36 with TFs like SIX homeobox 3b (six3b), Basic HLH (bhlhe23), Visual System Homeobox1 (vsx1), zinc finger homeobox protein 4 (zfhx4), D site albumin promoter binding protein b (dbpb), forkhead box G1b (foxg1b), basic helix-loop-helix family member e41 (bhlhe41) and inhibitor of DNA binding 2a (id2a) (Figure 3B). Each of these genes co-expressed with SMIM36 are developmental regulators of retinal gene expression and several are linked to human diseases. Out of the above TFs, six3b, with the highest co-expression index with SMIM36, is a well-established master regulator of forebrain and eye development (Inbal et al. 2007; Pineda-Alvarez et al. 2011), with loss of function of six3b leading to the absence of optic structures and ectopic expression leading to formation of eye like structures. The role of bhlhe23 in the formation of bipolar cells is known (Dong et al. 2020a; Woods et al. 2018; Dong et al. 2020b)), also vsx1 serves as a well-known marker of bipolar cells (Ohtoshi et al. 2004; Shi et al. 2011; Chow et al. 2001). Overall, co-expression analysis revealed the strong regulatory role of bipolar cell specific TFs in SMIM36 regulation. Additionally, reactome pathway analysis of top one percentile target genes of these TFs showed that SMIM36 is co-expressed with other retinal genes like snap25b, fam107b and rgs16 (Figure 3C).

**Figure 3.**
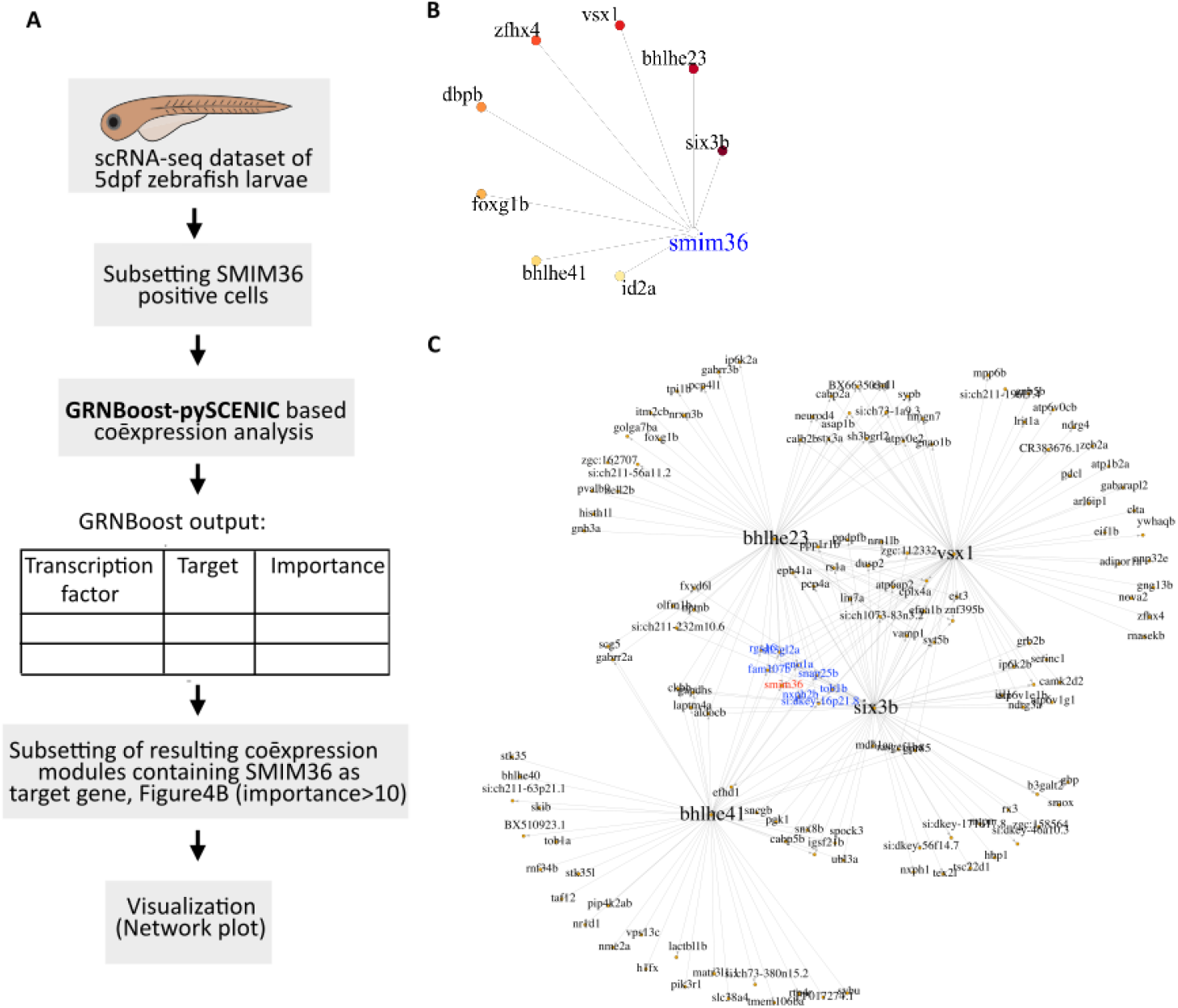
Co-expression analysis reveals potential regulation of SMIM36 by bipolar cell specific transcription factors. A) Schematic depicting workflow of the co-expression analysis using pySCENIC. B) Network plot showing the co-expression of SMIM36 with transcription factors (importance score>10), edges of the network plot are placed from highest (red) to lowest (yellow) importance score. C) Network plot depicting co-expression of top one percentile target genes of the transcription factors derived from SMIM36 module (Figure 3B) namely six3b, bhlhe23, vsx1 and bhlhe41. Gene MANIA reported co-expression networks are highlighted in blue color.

### Knockdown of SMIM36 affects eye size and retinal lamination in the zebrafish retina

The above findings highlight SMIM36 as a conserved microprotein across vertebrates. Additionally, its specific expression in the retinal bipolar cells and co-expression with master regulators of eye development suggest its probable function in a cell-type specific manner. To understand the in vivo function of SMIM36 in retinal physiology, we again turned to the vertebrate model, zebrafish. The loss of function of SMIM36 was achieved by a knockdown approach using an antisense morpholino. In zebrafish, SMIM36 contains two exons and a single intron; we designed antisense morpholinos targeting the splice junction of the intron and exon2. The microinjection of splice block morpholinos in single-cell zebrafish embryos led to the downregulation of SMIM36 transcript expression (Figure 4A). We followed the development of morphant larvae and observed larval death beyond the 7dpf stage, suggesting an indispensable role of SMIM36 in development. Since retinal development is completed before this time, all further experiments were conducted in 2 to 5 dpf larvae.

**Figure 4.**
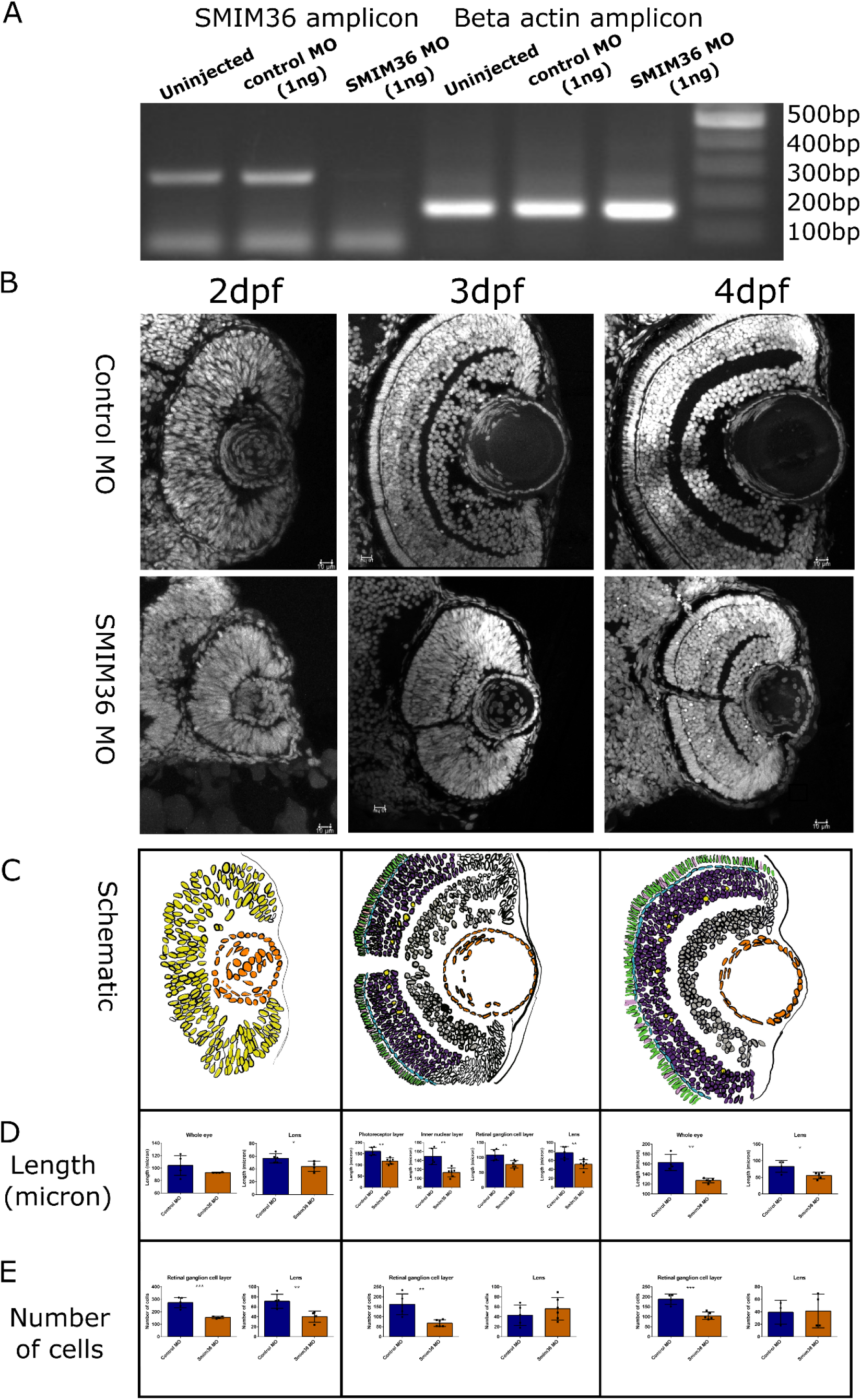
SMIM36 knockdown leads to microphthalmia and lamination defects in the zebrafish retina. A) RT-PCR validation of SMIM36 knockdown. gel image showing SMIM36 and beta actin amplicon in uninjected, control MO and SMIM36 MO injected zebrafish larvae (3dpf). B) Fluorescence images of hoechst stained retinal sections of zebrafish larvae injected with ControlMO and SMIM36 MO at 2dpf, 3dpf and 4dpf. scale bar-10μm, section thickness-10μm. C) Schematics of control retinal sections at 2dpf, 3dpf and 4dpf showing the retinal cell layers in different colors. Photoreceptor layer (green- cones, pink- rods), horizontal cells - blue, inner nuclear layer (purple-muller glia, yellow-bipolar cells), retinal ganglion cell layer- gray, lens cells – orange D) Quantification of the length of different cell layers in 2dpf, 3dpf and 4dpf retinal sections. E) Quantification of the number of cells in selected cell layers in 2dpf, 3dpf and 4dpf retinal sections.

The knockdown of SMIM36 caused a reduction in eye size (Figure 4B) of the zebrafish larvae. Further, a deeper look at retinal layers revealed retinal lamination defects in larvae injected with SMIM36 morpholino as compared to control morpholino (Figure 4B-E). The cells were arranged in well organised layers in the retina of wild type larvae (5dpf) but in the morphant, all the cells were in the same layer. Further, total cell counts were similar; the cells were not arranged in the radial manner and they also showed reduced polarisation. The inner plexiform layer and outer plexiform layers which separate the nuclear layers were absent. Similarly, the highly polar photoreceptor cell layer was also absent in the morphant.

To establish that the protein coding region of the SMIM36 gene is sufficient to carry out its function, we microinjected in vitro synthesised transcripts containing the SMIM36 coding sequence (CDS) along with the morpholino injected embryos which led to the partial rescue of knockdown mediated phenotype, suggesting the specificity of the observed phenotype (Figure 5A-C). Although the eye size was partially rescued, closer examination of the retinal layers showed that partial restoration of SMIM36 expression had different effects in the various retinal layers. It was unable to rescue the photoreceptor layer but reduced the crowding of cells in the outer nuclear layer. Next, we mutated the SMIM36 coding region in an attempt to identify the domains that are necessary for its proper function. The domain structure of SMIM36 (Figure1A) shows that the single transmembrane domain (residues) separates a 10aa long N-terminal projecting outside and a 11aa long C-terminal domain that would project into the lumen. We created mutants lacking the first 51 and 37 amino acids named as Δ51 and Δ37 respectively, deleting the N-terminal end and the transmembrane domain. Microinjection of RNAs coding for truncated forms of SMIM36 did not rescue the morpholino mediated phenotype, hinting towards the essential role of the full length peptide (Figure 5D).

**Figure 5.**
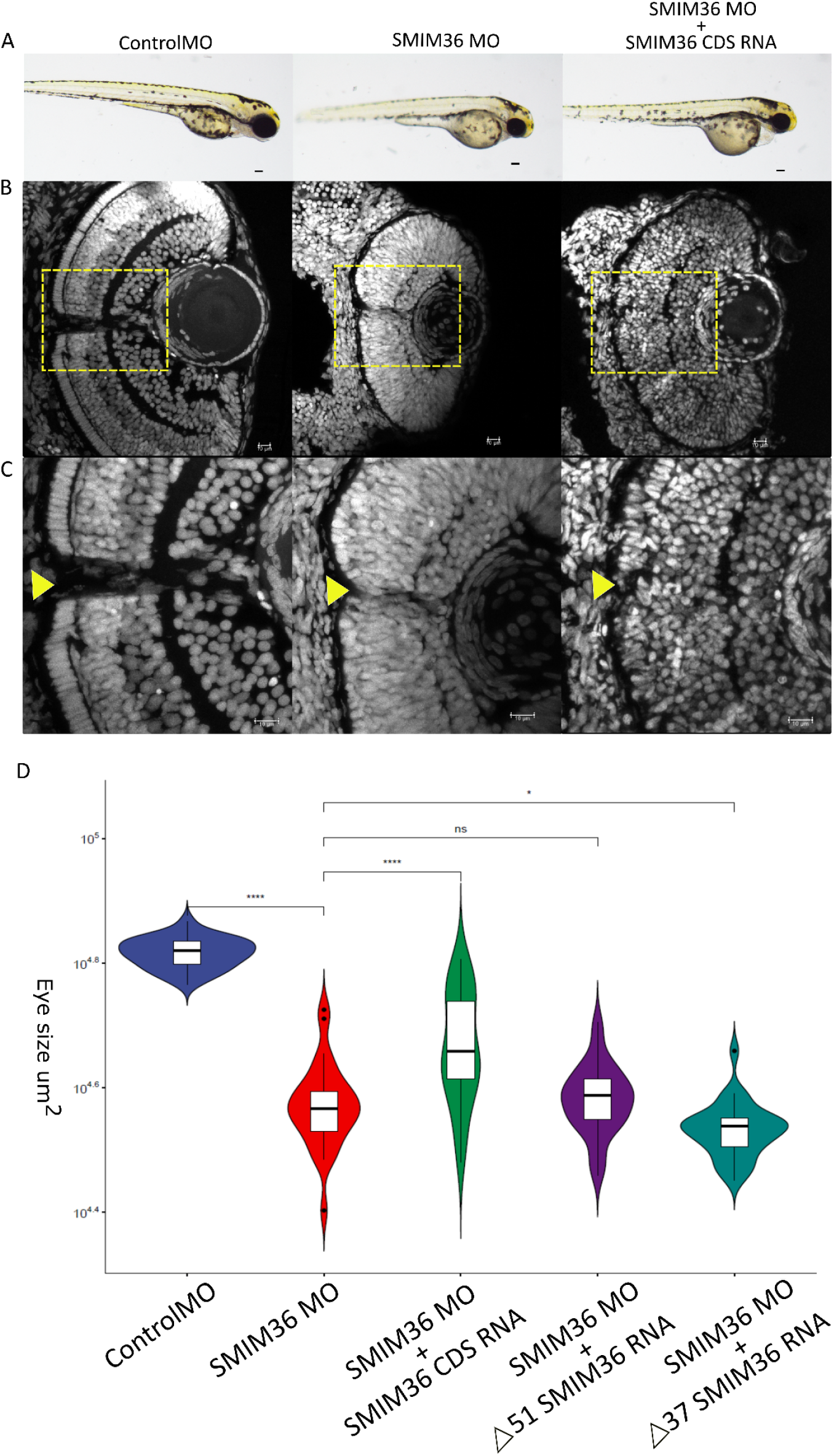
SMIM36 knockdown defects are partially rescued by full length SMIM36 peptide. A) Bright field images showing lateral view of zebrafish larvae injected with ControlMO, SMIM36 MO and SMIM36 MO+SMIM36 CDS RNA (rescue) at 3dpf (days post fertilization), scale bar-100μm. B) & C) Fluorescence images of hoechst stained retinal sections of zebrafish larvae injected with ControlMO, SMIM36 MO and SMIM36 MO+SMIM36 CDS RNA at 3dpf, the yellow arrowhead indicates optic nerve exit. Inset showing enlarged view of retinal layers, scale bar- 10μm, section thickness-10μm. D) Violin plot showing eye size measurement of zebrafish larvae injected with ControlMO, SMIM36MO and SMIM36MO with SMIM36 CDS RNA, Δ51SMIM36 RNA, Δ37 SMIM36 RNA at 3dpf, eye size calculated by manually drawing a circle around eye in ImageJ, \*\*\*\**p*<0.001, \**p*<0.05, ns -not significant.

Retinal development occurs in a coordinated fashion where each cell type is born in a specified temporal window. The later stages of retinal development are characterised by the migration of cell types to their designated layers. In the zebrafish retina, by 3dpf, all retinal layers are visible with functional synapses formed among cell types. By 5dpf of their development, zebrafish larvae show functional visual response as evidenced by 100% optokinetic response by 5dpf (Morris and Fadool 2005). Considering the cell-type specific expression of SMIM36 (Figure 6A), we hypothesised that retinal lamination defects are due to the aberrant formation of the particular cell type. We assessed the status of various cell types in the morphant retina, by performing qPCR for marker genes of other retinal cell types in the eyes of 5dpf morphant larvae (Figure 6B-H). In spite of the crowding of cells in the outer nuclear layer, the expression of the bipolar cell marker (prkcaa) showed a clear band in the morphant, not very different from the wildtype (Figure 6I). The markers for Muller glia (*glusyn)* and horizontal cells (*cx55*.*5)* were specifically decreased while all the other markers for cell types remained unaffected. This agrees well with the absence of clear plexiform layers because the Muller glia and horizontal cells are found in this layer. Thus, we found that the loss of SMIM36 resulted in selective loss of certain cell types and corresponding retinal layers leading to an overall reduction in eye size.

**Figure 6.**
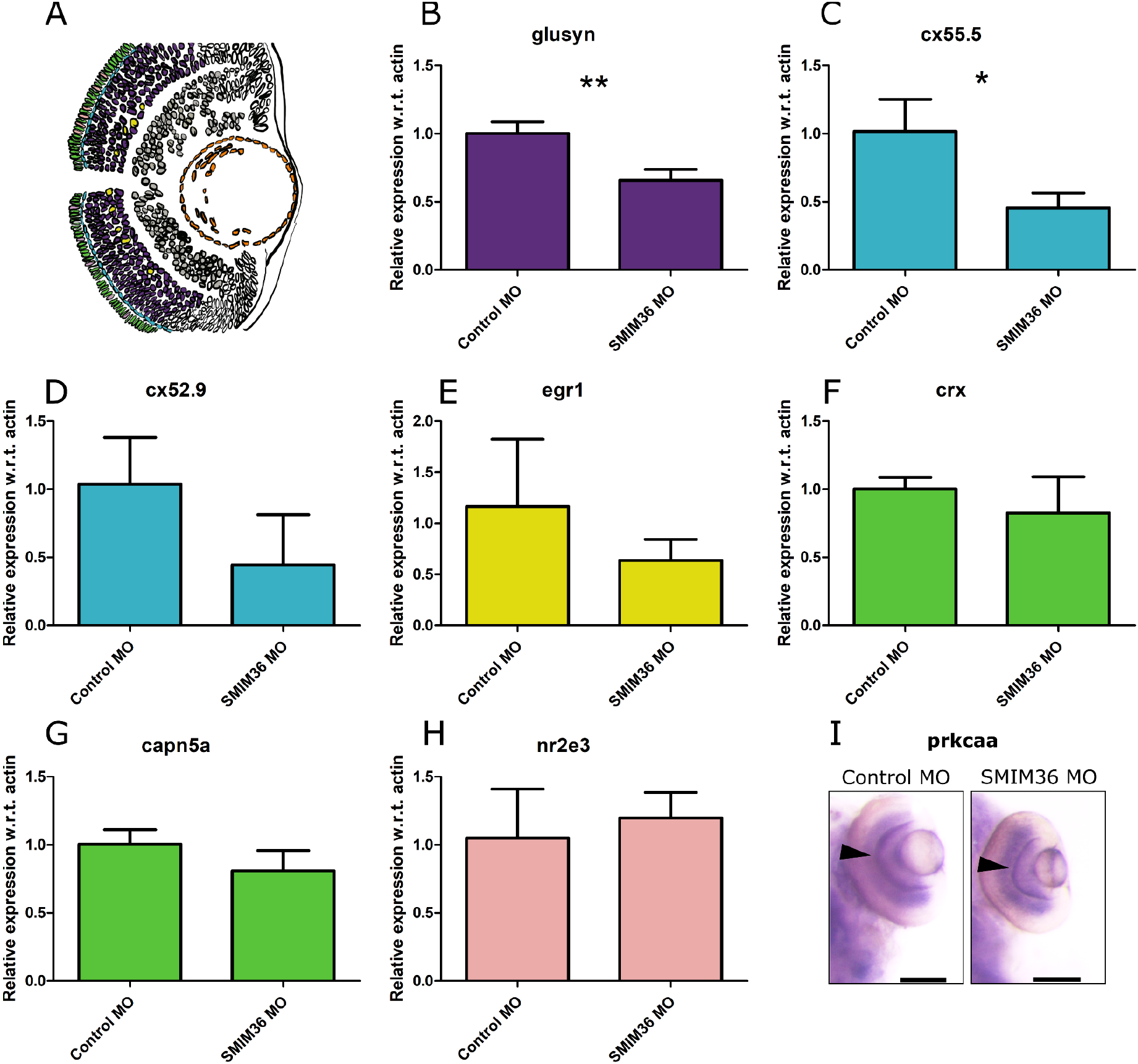
SMIM36 knockdown affects the mRNA expression of markers for Muller glia and horizontal cells. A) A schematic of the zebrafish eye section marking cell types with different colors. green-cones, pink-rods, blue-horizontal cells, purple-muller glia, yellow-amacrine cells. B) to H) Expression quantification of markers of retinal cell types in the eye of 5dpf zebrafish larvae injected with Control MO and SMIM36 MO by qRT-PCR. Bars represent mean±s.e.m. across three biological replicates (with ∼40 eyes per replicate), eyes from 5dpf control and morphant larvae were dissected and used for RNA extraction \*\**p*<0.01, \**p*<0.05. Retinal cell markers-Muller glia (glusyn), horizontal cells (cx55.5, cx52.9), amacrine cells (egr1), cones (crx, capn5a),rods (nr2e3). I) Whole mount In situ hybridization of bipolar cell marker, *prkcaa* in 5dpf larvae injected with ControlMO and SMIM36 MO, scale bar-100μm.

## Discussion

The zebrafish retina is anatomically and functionally similar to the human retina, thus presenting a model to study retinal degeneration (Angueyra and Kindt 2018). Due to external fertilization followed by rapid development, within 72 hours post-fertilization, all the structures of the adult retina are formed and easily visualized. For instance the first ganglion cells exit the cell cycle and start differentiation by 20 weeks of gestation in humans, but can be seen in zebrafish within 40 hours post fertilization (Richardson et al. 2016). Zebrafish mutants recapitulate the defects seen in a number of human ocular conditions like coloboma, cataract and cyclopia. Loss of SMIM36 resulted in microphthalmia, previously shown by mutants in GDF6, PAX6 and SIX6. Loss of Six3b, the gene that is most highly co-expressed with SMIM36 also shows microphthalmia(Inbal et al. 2007). Zebrafish is also amenable to chemical screens to identify small molecules that can compensate for the developmental defects of mutants.

We explored GWAS data to check if the SMIM36 locus is implicated in any eye diseases. Interestingly, the human SMIM36 gene lies within a region implicated in High Myopia, also known as degenerative myopia, a genetic condition characterized by extreme ‘nearsightedness’ and a pre-disposition to retinal detachment. Further studies are required to explore mutations in the human SMIM36 gene. The availability of a zebrafish model can be useful in screening for mutations, drug screening and in depth functional studies.

SMIM36 codes for a small peptide belonging to a class of microproteins. Microproteins are recently annotated and their importance in diverse cellular functions is being unravelled rapidly. To the best of our knowledge, there does not exist any study mentioning the function of microproteins in the neural system. Often, miroproteins arise from non-canonical loci in the genome like previously annotated non-coding regions and the 5’ and 3’ UTR region of well-known large ORFs (Chen et al. 2020). The conservation in the protein sequence of SMIM36 points to a vital function of this novel microprotein and its highly restricted expression pattern suggests that it is involved in the development or functioning of the retina. SMIM36 is expressed in the bipolar cells of zebrafish and human retina, with human retina showing expression in Muller glia and rods too.

The gene structure of SMIM36 shows striking similarity and significant differences between human and zebrafish. The untranslated regions of the mRNA SMIM36 are exceptionally long in both zebrafish and humans, while the small coding sequence arising from exon1 in both species is highly conserved. In zebrafish, the SMIM36 mRNA contains 484nt long 3’UTR whereas the human counterpart has 2397nt long 3’UTR. A number of studies have shown an increase in the length of 3’UTR in higher organisms (Wang et al. 2019; Xiong et al. 2018), thus enhancing the chances of fine regulation by affecting the binding of RNA binding proteins and miRNAs. We speculate that the long UTR region could be involved in the regulation of SMIM36, especially its rhythmic expression, an interesting aspect to explore in the future.

The loss of SMIM36 in zebrafish results in a very specific defect, the loss of Muller glia and horizontal cells. It is known that during retina development the migration and positioning of cell types are highly dependent on intercellular cues (D’Souza and Lang 2020). It is important to note that the expression of SMIM36 is not restricted to the cells that are affected by its loss.

SMIM36 protein might be assisting in cell-cell communication in the retina..To the best of our knowledge, no microprotein has so far been implicated in cell-cell interactions. However, being a transmembrane protein, SMIM36 expressed in the highly laminar layers of the retina, it is in a position to engage in local interactions between cells This is further strengthened by the retinal lamination defects observed in SMIM36 morphant retina suggesting that loss of expression in one cell type might have led to the disordered arrangement of neighbouring cellsIn the future, exploring protein-protein interaction partners would reveal the molecular mechanism of SMIM36.

In summary, we report that the zebrafish SMIM36 gene encodes a microprotein that is required for the proper development of the retina, specifically for the differentiation and laminar arrangement of the retinal layers. Our study paves the way for deeper investigations into the molecular function of this microprotein, chemical screening for small molecules and identifies SMIM36 as a candidate for human genetics of ocular diseases marked by the loss of Muller glia and horizontal cells.

## Materials and Methods

### Animal husbandry

Zebrafish were raised and bred according to standard protocols as described in the Zebrafish book (Westerfield 2007). The handling of zebrafish and experiments were performed in accordance with protocols approved by the Institutional Animal Ethics Committee (IAEC) of the CSIR-Institute of Genomics and Integrative Biology, India.

### Cloning, transfection and immunolabeling

A plasmid for mammalian expression of SMIM36 under CMV promoter was generated in a backbone of pEGFP-C1 vector. The sequence of human SMIM36 was obtained by gene synthesis in plasmid backbone (Genescript). The cloning cassette consisting of the SMIM36 sequence was amplified using primers flanking the CDS. The HindIII and Sal1 restriction sites were added in the forward and reverse primers respectively. The PCR product was cloned into the pEGFP-C1 vector digested with HindIII and SalI restriction enzymes using T4DNA ligase.

HeLa cell line was cultured in DMEM media (Invitrogen) supplemented with 10% FBS (Invitrogen) at 37°C in humidified chamber containing 5% CO_2_. 500 ng of pEGFP-C1 and pEGFPC1-SMIM36 plasmids were transfected in cells grown on four chambered cover glass using lipofectamine. After 24 Hours cells were fixed in 4%PFA for immunolabeling.

The immunolabeling was performed using a standard protocol, primary antibody for EGFP (ab290, 1:500) was incubated overnight at 4°C in a humid chamber. The secondary antibody was added at 1:1000 dilution in the blocking buffer for 1 hour at room temperature. DAPI was used for staining the nuclei, imaging was performed with 63x objective in Leica SP8 confocal microscope.

### RNA extraction and quantitative reverse-transcription PCR (qRT-PCR)

Total RNA from zebrafish larvae was extracted using trizol, the 500 ng RNA was used for making cDNA (QuantiTect Reverse Transcription Kit), 1μl of cDNA was used in the qPCR reaction. qPCR was performed using SYBR green PCR mix (TAKARA) in Roche Light Cycler.

### Single-cell RNA-seq (scRNA-seq) and co-expression network analysis

For human retina the scRNA-Seq dataset (n=3, Lukowski et al. 2019) was downloaded from (https://www.ebi.ac.uk/gxa/sc/experiments/E-MTAB-7316). The count data was then analysed in Seurat v3.1.5, the quality check and dimension reduction was performed using the same parameters as Lukowski et al. The subset analysis was performed by extracting the raw UMI matrix of SMIM36 expressing cells from total cells and subjected to Seurat analysis at a resolution of 0.6 with 5 principal components used for clustering.

For co-expression network analysis, the scRNA-seq dataset of (Spanjaard et al. 2018) was analyzed in Seurat. The count data of the SMIM36 expressing cluster was extracted and subjected to GRN-Boost in the pySCENIC pipeline for co-expression analysis. The GRN-Boost output file contains a list of transcription factors (TFs), target genes and their importance score. The TF-target gene pairs with an importance score of >10 were plotted in the network plot.

### Microinjection

Single-cell zebrafish embryos were injected with 1ng of splice block morpholino (MO) targeting the SMIM36, for standard control MO from Gene tools was used as control. For the rescue experiment, 350pg of RNA containing the coding sequence (CDS) of SMIM36 was injected. The RNA was in vitro transcribed (IVT) by mMessage mMachine T7 ultra kit (Invitrogen), followed by polyA tailing (Invitrogen). The IVT template was made by touchdown PCR using plasmid (containing SMIM36 CDS) as template. The RNAs for various truncated form of SMIM36 were also prepared following the same protocol.

Primer sequences used in the work are provided in the supplementary table.S1

### Imaging

The bright field imaging of 3dpf larvae was performed at 5x magnification, the eye size was calculated by manually drawing a circle around the eye for each image in imageJ. The hoechst stained fluorescent images of retinal sections were captured in Leica TCS SP8 confocal microscope using oil immersion 63x objective.

### Whole mount *In situ* hybridization

The plasmids for preparing *in situ* probe of *prkcaa* were kindly gifted by Dr. Stephen C. Neuhauss (University of Zurich, Switzerland) (Haug et al. 2018). 1μg linearized plasmid was used for In Vitro transcription to make DIG labelled RNA probe (DIG RNA Labeling Mix-11277073910 Roche). Whole mount RNA *in situ* hybridization was performed by following the standard zebrafish protocol (http://zfin.org/ZFIN/Methods/ThisseProtocol).

## Acknowledgements

We acknowledge the MLP2102 project for funding this work and CSIR for providing student fellowship. We also acknowledge the zebrafish facility, imaging facility and high computing facility of CSIR-IGIB. We acknowledge Dr Souvik Maiti, CSIR-IGIB for providing funding and inputs during discussions for proceeding the work.

## Author contributions

BP and SS designed the experiments, SS performed the zebrafish experiments, imaging and scRNA-seq analysis, SR performed the co-expression analysis, YK did cell line transfections and mammalian expression vector cloning, DCJ did the validation of zebrafish experiments, BP and SS wrote the article.

